# Does linked selection explain the narrow range of genetic diversity across species?

**DOI:** 10.1101/042598

**Authors:** Graham Coop

## Abstract

The relatively narrow range of genetic polymorphism levels across species has been a major source of debate since the inception of molecular population genetics. Recently Corbett-Detig et al found evidence that linked selection strongly constrains levels of polymorphism in species with large census sizes. Here I reexamine this claim and find weak support for this conclusion. While linked selection is an important determinant of polymorphism levels along the genome in many species, we currently lack compelling evidence that it is a major determinant of polymorphism levels among obligately sexual species.

---

Two observations have puzzled population geneticists since the inception of molecular population genetics. The first is the relatively high level of genetic variation observed in most obligately sexual species. The neutral theory of molecular evolution was developed in part to explain the high levels of diversity. Under a simple neutral model, with constant population size, we should expect the amount of neutral genetic diversity to scale with the product of the population size and mutation rate. Specifically, in a randomly mating diploid population of size *N*_*c*_ individuals, at sites that experience a neutral per generation mutation rate of μ, we should expect pairwise diversity *π* ≈ 4*N*_*c*_*μ*. The second observation, however, is the relatively narrow range of polymorphism across species with vastly different census sizes (see Leffler *et al*., 2012, for a recent review). As highlighted by Lewontin (1974) in his discussion of the paradox of variation, this seemingly contradicts the prediction of the neutral theory that genetic diversity should scale with the census population size. In eukaryotes, per generation mutation rates appear to vary over just two orders of magnitude (10^−10^–10^−8^), while census population sizes range over many orders of magnitude, from populations consisting of just a few thousand individuals to populations measured in billions of individuals. Therefore, diversity levels should vary over many orders of magnitude, but levels of synonymous genetic diversity in obligately sexual populations differ by only a few, from 0.01 – 10% (Leffler *et al*., 2012).

There are a number of explanations for the discrepancy between genetic diversity levels and census population sizes. The first is that the effective size of the population (*N*_*e*_) is often much lower than the census size, due to high variance in reproductive success and frequent bottlenecks (see Charlesworth, 2009, for a review). The second major explanation, put forward by Maynard Smith and Haigh (1974), is that neutral levels of diversity are also systematically reduced by the effects of linked selection. The sweep of a beneficial allele to fixation removes neutral diversity at linked sites (the hitchhiking effect; Maynard Smith and Haigh, 1974; Kaplan *et al*., 1989; Stephan *et al*., 1992) as does the removal of deleterious alleles from the population (background selection; Charlesworth *et al*., 1995; Hudson and Kaplan, 1995b,a; Nordborg *et al*., 1996). In fact, a wide range of models of selection predict the removal of neutral diversity linked to selected sites. This is because heritable variance in fitness leads the effect of high variance in reproductive success on levels of diversity to be compounded over generations (Robertson, 1961; Santiago and Caballero, 1995, 1998; Barton, 2000; Gillespie, 1994, 1997; Neher *et al*., 2013; Good *et al*., 2014).

Evidence for the action of linked selection in reducing levels of polymorphism includes a positive correlation between putatively neutral diversity and recombination in a number of species, as, all else being equal, linked selection should be less efficient in removing diversity in regions of high recombination (AguadÉ *et al*., 1989; Begun and Aquadro, 1992; Wiehe and Stephan, 1993; Cutter and Choi, 2010; Hellmann *et al*., 2008; Cai *et al*., 2009; Cutter and Payseur, 2013). In addition, levels of putatively neutral diversity are often lower in regions with a higher density of functional sites, where we expect more selected alleles to arise (Andolfatto, 2007; Macpherson *et al*., 2007; McVicker *et al*., 2009; Sattath *et al*., 2011; Hernandez *et al*., 2011; Beissinger *et al*., 2015). In large populations, selective sweeps and other forms of linked selection may come to dominate genetic drift as a source of stochasticity in allele frequencies, establishing an upper limit to levels of diversity (Kaplan *et al*., 1989; Gillespie, 2000).

To understand the contributions of the two explanations to levels of diversity, it is helpful to distinguish between the average observed level of neutral polymorphism in the genome (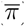) and that expected in the absence of linked selection (*π*_0_). Our idealized neutral level of variation in which *π*_0_ ≈ 4*N*_*e*_*μ*, reflects the effective size of the population (*N*_*e*_) in the absence of linked selection ( here *N*_*e*_ is not estimated simply from putatively neutral diversity levels genome-wide). To illustrate this point, take the extreme scenario in which linked selection explains nearly all of the deficit in variation in species with large census sizes, with fluctuations in population size playing a minor role. In these species, 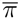 should be orders of magnitude smaller than *π*_0_, and *N*_*e*_ should be roughly the same order of magnitude as the census size. In contrast, if fluctuations in population size explain most of the deficit, then 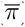 should be close to *π*_0_ for all species, while *N*_*e*_ would be many orders of magnitude lower than census population sizes for species with large population sizes.

Recently, Corbett-Detig *et al*. (2015) sought to test the idea that linked selection explains the narrow range of genetic diversity. They compiled published genomic data and genetic maps from 40 obligately sexual species to obtain an impressive dataset for comparative populations genomics. In these data, they found a positive correlation between diversity at four-fold degenerate sites and recombination rates in many species. To estimate the impact of linked selection, they then fit a simple model of hitchhiking and background selection to the relationship between recombination rate per base (*r*) and *π* in genomic windows within each species (also factoring in variation in gene density). Specifically, they fit a model where

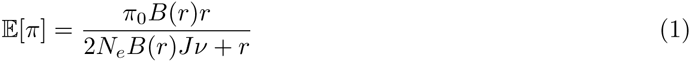

where *π*_0_ = 4*N*_*e*_*μ* is the expected diversity in the absence of linked selection, 1/*B*(*r*) is the increase in the rate of coalescence (genetic drift) due to background selection, *v* is the rate of sweeps per base pair, and *J* is related to the probability that a particular sweep forces a pair of lineages to coalesce averaged over the distance to the sweep (see Coop and Ralph, 2012, for more discussion). For full sweeps of an allele with an additive selection coefficient s, *J* ≈ *log*(2*N*_*e*_*s*)/*s*.

Corbett-Detig *et al*. (2015) found that their estimated reduction in diversity due to linked selection 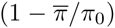 is positively correlated with measures of range size and inversely correlated with body size across species, both of which are correlates of the census population size (*N*_*c*_). The authors interpreted these findings as evidence that *“pervasive natural selection constrains neutral diversity and provides an explanation for why neutral diversity does not scale as expected with population size”*.

Stronger linked selection in species with larger population sizes is indeed qualitatively consistent with Maynard Smith and Haigh (1974)’s explanation of the narrow range of diversity (Kaplan *et al*., 1989). But the hypothesis that linked selection is the major contributor to the narrow range of diversity is a quantitative prediction and requires linked selection to dominate genetic drift by orders of magnitude in species with large population sizes (Kaplan *et al*., 1989; Gillespie, 2000). I present a reanalysis of the results of Corbett-Detig *et al*. (2015), focusing on these quantitative predictions.

Across the 40 species studied, the average level of diversity 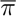 ranges from 0.1%–3.0% in animals and 0.012% – 1.5% in plants. The authors’ predictions of the range of diversity in the absence of linked selection (*π*_0_) are surprisingly similar (from 0.12%-3.36% in animals, and 0.012%–1.5% in plants). Even the maximum estimated reduction in genome-wide diversity (70%) is less than one order of magnitude. In order for linked selection to explain the paradox of variation, however, the estimated reductions would need to vary over several orders of magnitude to explain the reduction in variation in species over the full range of census population sizes.

To further illustrate this point, I plot the observed genome-wide average (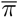) for each species versus the estimated level of diversity in the absence of linked selection (*π*_0_) (Figure 1). If linked selection strongly constrained levels of diversity, we should expect to see a pattern somewhat like the red line, in which species with very high levels of diversity in the absence of linked selection (*π*_0_) only have moderately higher levels of genome-wide average diversity (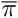). Instead, predicted diversities fall mostly along the *x* = *y* line — the result that we would expect if linked selection had only a limited impact on levels of diversity in large populations and diversity levels scaled with effective population sizes estimated in the absence of linked selection. This finding is consistent with the idea that demographic fluctuations are the principal determinant of levels of diversity among species. Interestingly, selfing species (shown in italics; Figure 1) seem to show the best evidence of large reductions in diversity due to linked selection, perhaps due to their much reduced effective rate of recombination (Nordborg, 1997; Charlesworth *et al*., 1997; Charlesworth and Wright, 2001).

**Figure 1:**
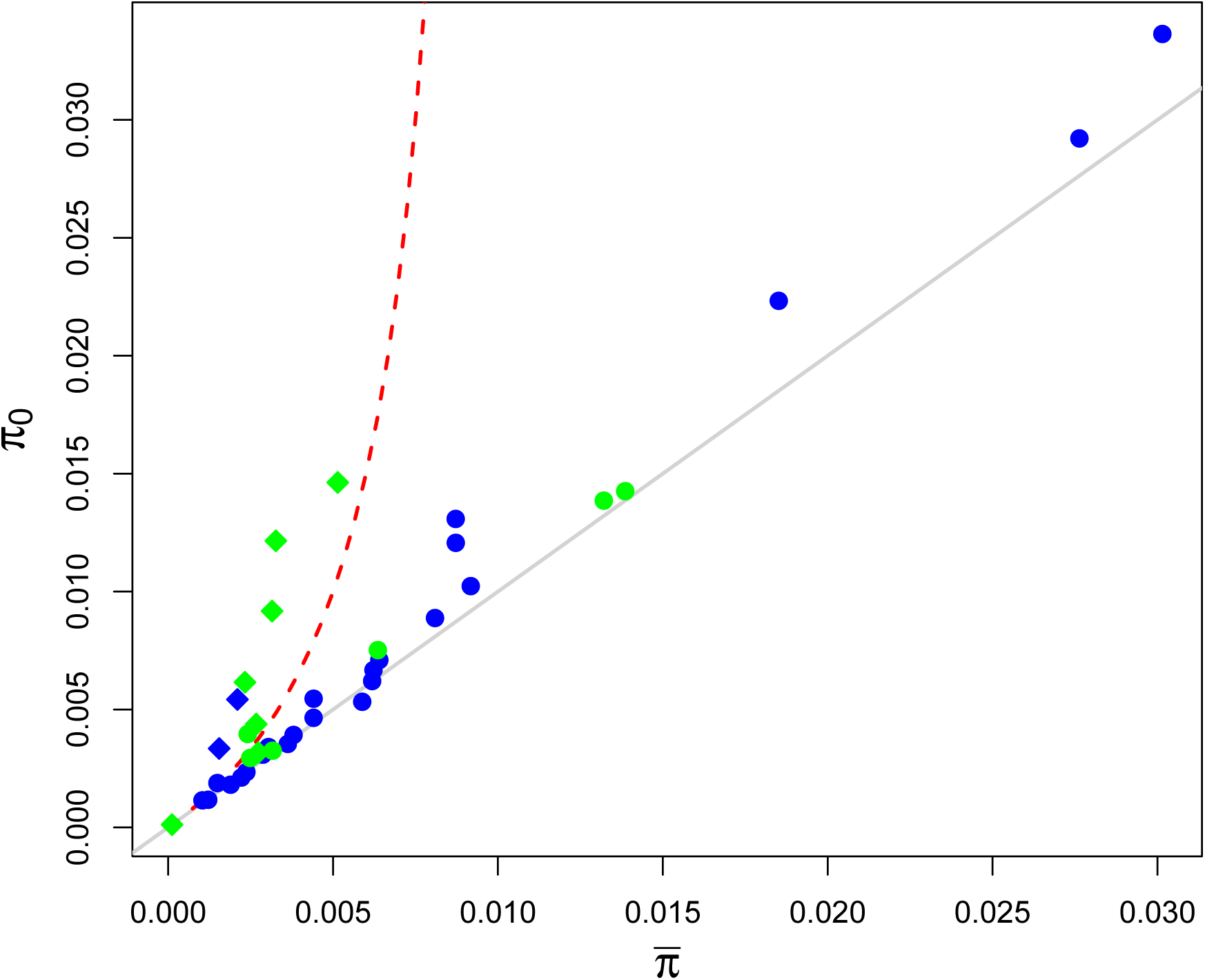
The genome-wide average of four-fold degenerate site diversity (*π*_*avg*_) plotted against Corbett-Detig *et al*. (2015)’s estimates of the level of genetic diversity in the absence of linked selection (*π*_0_) across species. Animal species are shown in blue, plant species are in green. Species with high selfing rates, as identified by Corbett-Detig *et al*. (2015), are shown as diamonds. The grey line shows the *x* = *y* line that we would expect in the absence of linked selection. The red line shows the type of relationship that we would expect to see if linked selection were a general explanation for the narrow range of genetic diversity. This example relationship was obtained by using a range of *N* in eqn. (1), setting *μ*_*BP*_ = 10^−9^, *r* = 10^−8^, and *v*_*BP*_ = 10^−11^ with *J* = 10^−4^ (roughly corresponding to a sweep of *s* = 10^−4^). I ignore background selection in this prediction for the sake of simplicity.

These results seemingly do not support the idea that linked selection dominates drift in the removal of diversity from large populations. That said, caution is warranted, as *π*_0_ was estimated by fitting a simple model of linked selection. In particular, these estimates assume that in regions of high recombination (and low enough gene density), we can hope to see close to *π*_0_, i.e. that the curve of *π* against *r* plateaus to *π*_0_ as drift comes to dominate linked selection. However, if linked selection truly dominates genetic drift, we may have little indication as to the true effective size of the population (i.e. in the absence of linked selection). For example, if the rate of coalescence due to hitchhiking, on the time-scale of genetic drift, is much higher than the highest recombination rate per base pair in the genome (2*N*_*e*_*vJ* ≫ *r*), then

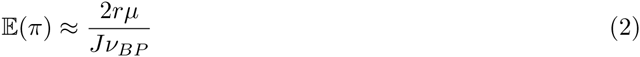

i.e. diversity increases linearly with recombination rate, and is completely independent of the rate of drift (Kaplan *et al*., 1989; Gillespie, 2000; Coop and Ralph, 2012).

To begin to explore whether levels of diversity in high recombination regions are still subject to strong linked selection or represent a balance of drift and mutation, in Figure 2 I plot the slope and correlation of pairwise diversity regressed against recombination both genome-wide and in windows of high recombination, using the binned data of Corbett-Detig *et al*. (2015). If linked selection dominated drift, we would expect to still see a positive slope between diversity and recombination even in high recombination regions. However, we fail to observe this pattern (right column of Figure 2) in the majority of species. The restricted recombination range in the second column means that we should expect the correlations to potential be lower, and the slope estimates to be noiser (i.e. have larger confidence intervals). However, the low upper confidence intervals for the slope in the top quartile of recombination regions suggest that the relationship between diversity and recombination is plateauing in most species. Thus there is preliminary evidence that recombination — and thus presumably linked selection — is not a strong determinant of levels of diversity in high recombination regions.

**Figure 2:**
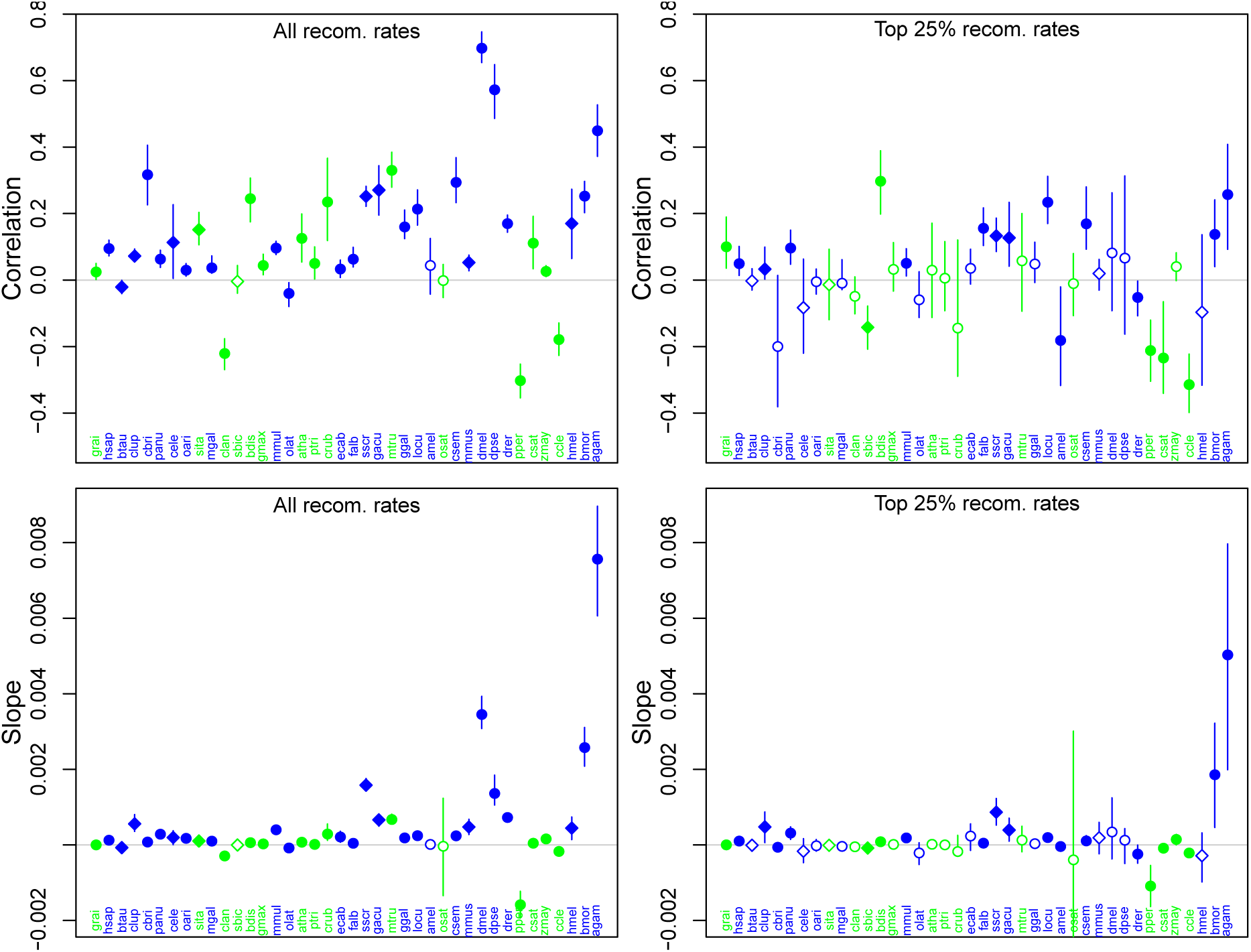
**Top row** the Pearson correlation coefficient between recombination rate and genetic diversity at fourfold degenerate sites in 500kb windows, colored and labeled by species. On the left, all windows are used, on the right only windows with recombination rates in the top quartile. Species are arranged left to right in order of increasing average levels of diversity. Points are colored as in Figure 1. The line shows the 95% bootstrap confidence interval for the correlation coefficient calculated by sampling 500kb windows with replacement. Open points indicate that the 95% confidence interval overlaps zero. **Bottom row** the slopes of the linear regression of diversity on recombination rate (with the same layout as the top panel).

These findings suggest that, in many species, levels of diversity in high recombination regions potentially do offer good estimates of the effective population size in the absence of linked selection. Thus, even in the apparent absence of linked selection, the range of effective population sizes appears tightly constrained across species that differ vastly in their census sizes.

This is not to say that linked selection is not an important factor in understanding levels of diversity within genomes. Undoubtedly as genetic maps, functional annotations, and statistical methodologies improve, our appreciation of the effects of linked selection will increase as well (McVicker *et al*., 2009; Comeron, 2014; Elyashiv *et al*., 2014). For example, Elyashiv *et al*. (2014) inferred an average diversity reduction of only 34-36% in *Drosophila melanogaster* using just the relationship between recombination rates and diversity, similar to that used here, and found no effect of linked selection in the upper 1% quantile of predicted diversity levels. However, by fitting a more detailed model that incorporated a range of annotations and variation in the strength of selection, Elyashiv *et al*. (2014) estimated an average 77 - 89% reduction in diversity due to selection on linked sites, and concluded that no genomic window was entirely free of the effect of selection. Nonetheless, it seems unlikely that our estimates of the effect of linked selection (based on current models) will approach the orders of magnitude needed for linked selection to be the major explanation of the paradox of variation.

In summary, while we still lack a full picture, a reanalysis of the results of Corbett-Detig *et al*. (2015) does not support the idea that linked selection is the major factor in shaping diversity levels among obligate sexual species. As highlighted by Corbett-Detig *et al*. (2015), linked selection does play a major role in shaping variation levels *along* many species’ genomes, sometimes contributing more to variation in allele frequencies than genetic drift alone. As such, Corbett-Detig *et al*. (2015)’s results reinforce the view that linked selection must be more thoroughly incorporated into our null models for population genetics. But it may well be the case that the major driver of variation in levels of diversity *among* sexual species, and therefore the major cause of the discrepancy between levels of diversity and population size, is fluctuations in population size.

## Acknowledgements

I thank Jeremy Berg, Gideon Bradburd, Vince Buffalo, Nancy Chen, Chuck Langley, Molly Przeworski, Jeff Ross-Ibarra, and Guy Sella for useful discussions based on earlier drafts of this note. I thank Russ Corbett-Detig and Tim Sackton for their feedback on an earlier draft and for making their code and data publicly available.

